# An efficient and stable ascorbate/O_2_‑driven route for L‑DOPA synthesis by heme‑dependent tyrosine hydroxylase

**DOI:** 10.64898/2026.04.24.720734

**Authors:** Langxing Liao, Zongyu Bao, Zhihui Jiang, Aitao Li, Binju Wang

**Author notes:** These authors contributed equally to this work (Langxing Liao, Zongyu Bao, Zhihui Jiang).

## Abstract

L-DOPA is a key therapeutic agent for Parkinson’s disease, with growing demand due to global population aging. Here we report that heme-dependent tyrosine hydroxylase (TyrH) can utilize an ascorbate/O_2_ system—as an alternative to H_2_O_2_—to synthesize L-DOPA with markedly enhanced operational stability. While exogenous H_2_O_2_ rapidly inactivates TyrH within minutes, sodium ascorbate (NaAsc) enables sustained catalysis for up to 24 h, surpassing the H_2_O_2_-driven yield after only 30 min. UV-vis spectroscopy confirms that H_2_O_2_ readily degrades the heme center, whereas the heme remains intact in the presence of NaAsc. QM/MM simulations reveal that *in situ* generated H_2_O_2_ leads to the active species of Compound I for tyrosine hydroxylation. Through systematic optimization, we establish efficient reaction conditions (40 µM TyrH, 1 mM L-Tyr, 100 mM NaAsc, pH 8.5, 40 °C), achieving >95% conversion of L-Tyr to L-DOPA within 2 h. This work not only provides a robust and sustainable biocatalytic route for L-DOPA production but also highlights the broader applicability of the ascorbate/O_2_ pathway in heme-enzyme catalysis.

## Introduction

Since the 1960s, levodopa (3,4-dihydroxy-L-phenylalanine, L-DOPA) has been the primary treatment for Parkinson’s disease, a neurodegenerative disorder caused by dopamine depletion in the brain.^1, 2^ Unlike dopamine, L-DOPA crosses the blood–brain barrier and effectively elevates dopamine levels in patients.^3, 4^ To this day, L-DOPA remains the first-line therapy for Parkinson’s disease, and its demand continues to rise with global aging trends.^5^ Currently, L-DOPA production relies mainly on asymmetric chemical synthesis,^6^ which suffers from low conversion efficiency and poor enantioselectivity.^7^ In contrast, biosynthesis offers advantages such as mild reaction conditions, simplified procedures, and environmental compatibility.^8^ Through metabolic engineering, microorganisms can be designed to heterologously express key enzymes—including tyrosinase,^9-11^ *p*-hydroxyphenylacetate 3-hydroxylase (PHAH),^12-14^ tyrosine phenol-lyase (TPL),^15-17^ or L-tyrosine hydroxylase (TyrH)^18, 19^—to serve as biocatalysts for L-DOPA production. These biotechnological approaches improve conversion efficiency, yield, enantioselectivity, and overall cost-effectiveness.^8^

Conventional TyrH is a pterin-dependent mononuclear non-heme iron oxygenase that catalyzes the hydroxylation of L-tyrosine (L-Tyr) to L-DOPA in catecholamine neurotransmitter biosynthesis.^18^ A distinct group of microbial enzymes, the histidine-ligated heme-dependent TyrHs, initiate the biosynthesis of certain antibacterial and antitumor natural products in Actinomyces.^19-23^ These enzymes belong to the histidine-ligated heme-dependent aromatic oxygenase (HDAO) superfamily and can utilize H_2_O_2_ as an oxidant for L-Tyr hydroxylation.^24^ It has been reported that heme-dependent TyrH can also catalyze the reaction using an ascorbate (Asc)/O_2_ system, although its activity is two orders of magnitude lower than with H_2_O_2_.^22^ Experimental and computational studies indicate that Compound I (Cpd I) is generated by TyrH with H_2_O_2_, and this Cpd I subsequently mediates the hydroxylation of L-Tyr to L-DOPA.^22, 24-26^

Among heme-containing peroxidases, unspecific peroxygenases (UPOs) and P450 peroxidases can employ hydrogen peroxide as an oxidant to directly generate Cpd I, enabling oxygen incorporation into inert C–H bonds without complex electron-transfer chains.^27-39^ Similarly, copper-containing lytic polysaccharide monooxygenases can also employ H_2_O_2_ to generate high-valent metal-oxo species for the activation of C–H bonds.^40, 41^ However, exogenous H_2_O_2_ leads to irreversible enzyme inactivation, limiting industrial application._42_ Recently, these peroxidases were found to operate by pairing O_2_ with small-molecule reductants—such as Asc for UPOs and Asc, gallic acid or pyrogallol for LPMOs—to drive their respective oxidative reactions.^43-45^ Compared to the conventional H_2_O_2_-dependent pathway, the O_2_/small reductant-dependent pathway can mediate the substrate oxidation with high efficiency while avoiding the H_2_O_2_-induced inactivation of enzyme in the same time, highlighting its great potential for industrial applications.

In this study, we systematically gauged the catalytic behavior of TyrH with H_2_O_2_ or Asc/O_2_. We find that TyrH is rapidly inactivated during H_2_O_2_-driven reactions, severely restricting industrial utility. In contrast, the Asc/O_2_-dependent route exhibits quite high stability, with no observed inactivation of enzyme during catalysis. Furthermore, we systematically optimize the reaction conditions for TyrH-catalyzed conversion of L-Tyr for the sodium ascorbate (NaAsc)/O_2_-dependent system, identifying optimal parameters as 40 µM TyrH, 1 mM L-Tyr, 100 mM NaAsc, pH 8.5, and 40 °C. Under these conditions, near-complete substrate conversion is achieved with a catalytic efficiency of 500 µmol L^−1^ h^−1^, demonstrating its high potential for industrial application.

## Results and discussion

### Comparative Analysis of TyrH Activity and Stability in H_2_O_2_ versus NaAsc/O_2_ Systems

As reported previously,^22^ heme-dependent TyrH can hydroxylate L-Tyr to L-DOPA using either H_2_O_2_ or Asc/O_2_ systems (Fig. 1A). We first evaluated the activity of purified TyrH. HPLC analysis showed that TyrH produces L-DOPA from L-Tyr in the presence of H_2_O_2_, Asc, or NaAsc (Fig. 1B). We further analyzed the time course of TyrH-mediated hydroxylation using H_2_O_2_, NaAsc, or Asc (Fig. 1C). In the presence of H_2_O_2_, a high product concentration was observed within 1 min, followed by only a slight increase after 10 min. No further production was detected over time, indicating enzyme inactivation, with the L-DOPA concentration reaching only 7.3 μM. In contrast, with NaAsc, the reaction proceeded almost linearly over time, and the L-DOPA concentration surpassed that of the H_2_O_2_ system after 30 min, reaching 13.4 μM. After 2 h of reaction, the product concentration reached 51.8 μM. These results demonstrate that although TyrH exhibits a slower initial rate with NaAsc, it ultimately achieves a higher final yield, indicating better potential for practical application. When Asc was used, the L-DOPA concentration was comparable to that of the H_2_O_2_ system after 1 h, reaching 7.2 μM.

**Fig. 1.**
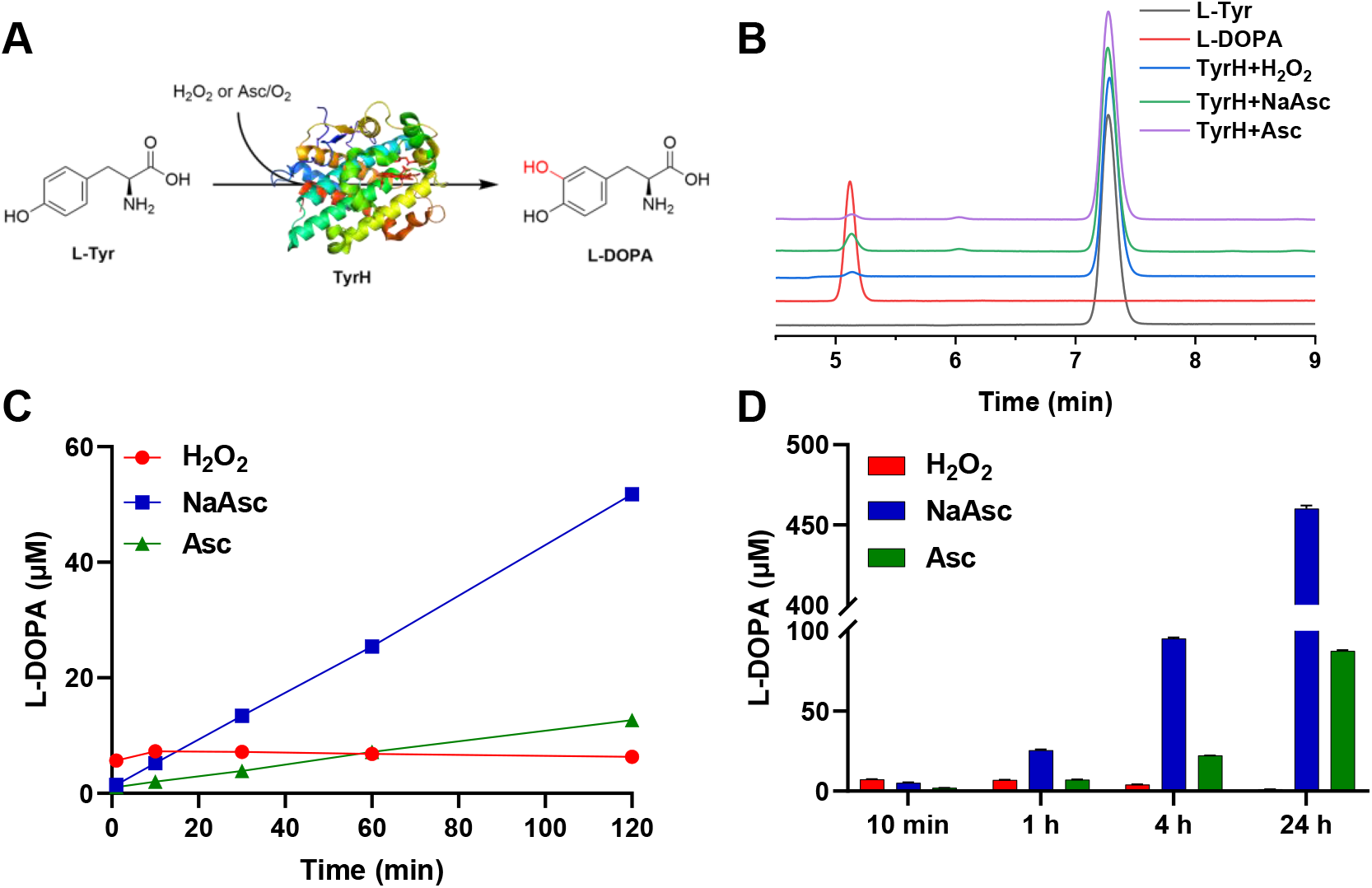
The hydroxylation of L-Tyr to generate L-DOPA by TyrH. (A) TyrH oxidizes L-Tyr to L-DOPA with either H_2_O_2_ or Asc/O_2_. (B) HPLC analysis of TyrH reaction with L-Tyr in the presence of H_2_O_2_, Asc, or NaAsc. (C) Time course of TyrH-mediated hydroxylation of L-Tyr with H_2_O_2_, Asc, or NaAsc in 2 hours. (D) The yield of L-DOPA produced by TyrH under H_2_O_2_, Asc, or NaAsc in different time. The standard reaction system contained 20 μM TyrH, 1 mM L-Tyr, and 1 mM H_2_O_2_ or 20 mM NaAsc/Asc at 30 °C in 50 mM Tris-HCl buffer, pH 7.5, 150 mM NaCl.

The highest L-DOPA concentration after 30 min was obtained with NaAsc, exceeding levels produced with H_2_O_2_ or Asc. The superior yield with NaAsc over Asc is attributed to the sharp pH drop caused by Asc addition, which limits L-DOPA production, whereas NaAsc does not alter the reaction pH. Product yields at different time points clearly indicated that NaAsc affords significantly higher yields than either H_2_O_2_ or Asc, reaching 460.2 μM after 24 h of reaction—in contrast to less than 100 μM for the other two systems—thereby confirming NaAsc as a more effective reductant for TyrH-mediated hydroxylation (Fig. 1D).

To gauge the TyrH stability with NaAsc or H_2_O_2_ (Fig. 2), we incubated the enzyme with either co-substrate and monitored UV-vis spectral changes. TyrH contains a histidine-ligated heme center with a characteristic Soret absorption peak at 403 nm for the resting enzyme, while absorbance strength at 403 nm reflects the concentration of the active TyrH. When TyrH was incubated with NaAsc, its absorption spectrum remained nearly unchanged even at higher NaAsc concentrations (Fig. 2A), indicating minor or no detrimental effect on TyrH activity. This is consistent with the linear L-DOPA production observed in time-course experiments with NaAsc (Fig. 1C). In contrast, H_2_O_2_ addition caused a rapid and pronounced decrease in the 403 nm peak (Fig. 2B), demonstrating heme degradation and concomitant enzyme inactivation, aligning with the loss of catalytic activity after ∼10 min in H_2_O_2_-driven reactions (Fig. 1C).

**Fig. 2.**
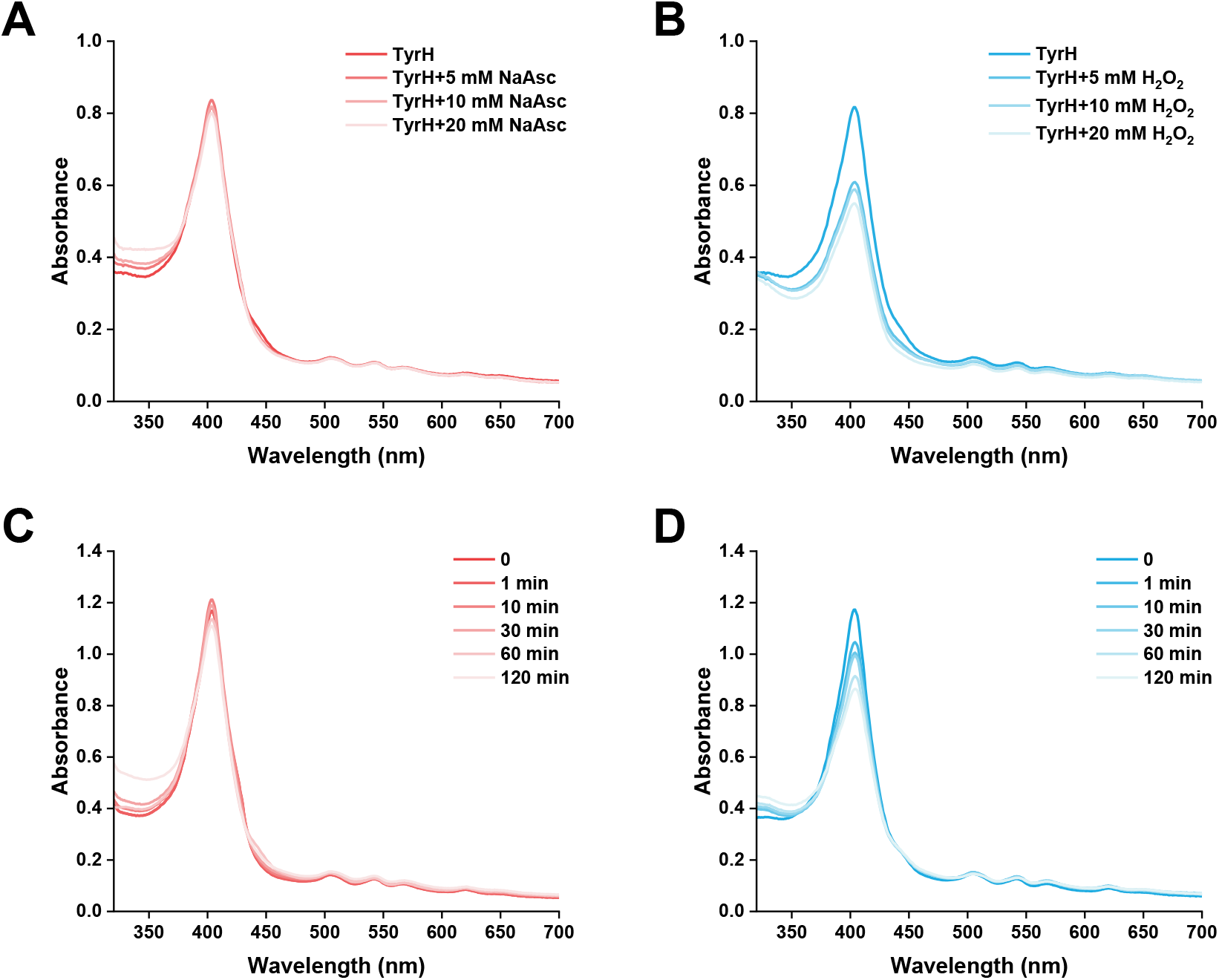
UV-vis spectra changes of TyrH treated with NaAsc or H_2_O_2_. The enzymes were treated with different concentrations of NaAsc (A) or H_2_O_2_ (B) in 50 mM Tris-HCl buffer, pH 7.5, 150 mM NaCl. The enzymes were treated with 20 mM NaAsc (C) or 1 mM H_2_O_2_ (D) in 50 mM Tris-HCl buffer, pH 7.5, 150 mM NaCl at different time. The treatment with H_2_O_2_ led to the pronounced heme-bleaching, while NaAsc treatment did not cause the heme-bleaching.

Under working concentrations, the 403 nm peak remained largely unchanged even after 2 h incubation with NaAsc, confirming the stability of the TyrH/NaAsc system for the sustainable applications (Fig. 2C). By contrast, the 403 nm peak drops rapidly within 1 min with H_2_O_2_ cosubstrate (Fig. 2D). The continued decline over time suggests progressive heme damage and irreversible loss of TyrH activity. Thus, UV-vis spectra reveal that H_2_O_2_ can induce the rapid inactivation of TyrH, while NaAsc is not.

### Mechanistic Investigation of TyrH with Ascorbate via QM/MM Calculations

Based on the high catalytic activity and stability of TyrH with NaAsc, we employed quantum mechanics/molecular mechanics (QM/MM) approach to elucidate the underlying reaction mechanism.

The QM/MM-calculated reaction pathway for O_2_ activation by TyrH is illustrated in Figure 3. Our QM/MM results reveal that the reactant complex of TyrH/Asc/O_2_ possesses an octuplet ground state (S = 7/2), in which two unpaired electrons are localized on the O_2_ moiety, while the remaining five unpaired electrons reside on the iron center. In the most favorable pathway, we found that the electron transfer (ET) from ascorbate to the ferric heme center is coupled to the O_2_ binding (^6^RC→^2^IM1), forming a Fe(III)–O_2_ ^−^ species within ^2^IM1(O–O distance, 1.29 Å). This process is calculated to be endothermic by 8.8 kcal/mol (^6^RC→^2^IM1). Starting from ^2^IM1, the hydrogen atom transfer from Ox of ASC radical to the distal oxygen of the Fe(III)–O_2_^−^ species is found to be quite facile (^2^IM1→^2^TS1), which involves a barrier of 0.8 kcal/mol and leads to the formation of Compound 0 within ^2^IM2. Starting from ^2^IM2, we found that a proton can transfer from a H88 residue to the proximal oxygen atom of Compound 0, resulting in the formation of a Fe(III)– H_2_O_2_ intermediate, which involves a barrier of 6.5 kcal mol^−1^ (^2^IM2→^2^TS2). Subsequently, the Fe(III)–H_2_O_2_ intermediate undergoes O–O bond heterolysis to generate the Cpd I intermediate, a process that is well studied in heme-dependent metalloenzymes.^46-48^ Finally, Compound I mediates the hydroxylation of L-Tyr, leading to the formation of the L-DOPA product.^25^

**Fig. 3.**
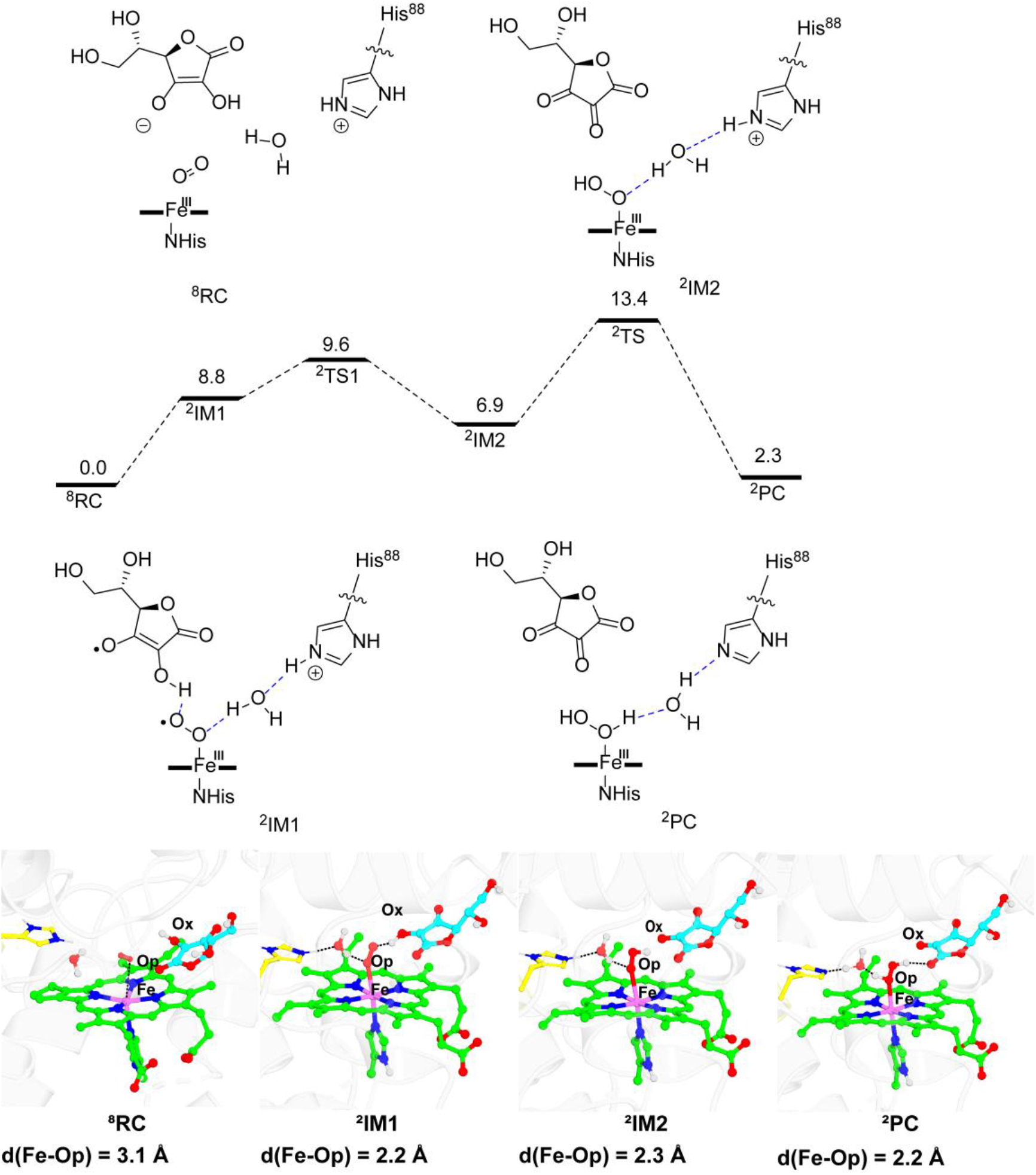
Mechanistic study of the O_2_ activation by TyrH in the presence of Asc. (A) QM(UTPSSH/def2-TZVP)/MM calculated mechanism and relative profile (in kcal mol^−1^) for Fe(III)–H_2_O_2_ formation from the TyrH/Asc/O_2_ system. (B) QM(UTPSSH/def2-SVP)/MM-optimized geometries of key species involved in the reaction.

### Systematic Optimization of Reaction Conditions for TyrH Using NaAsc

Although TyrH exhibits superior stability with NaAsc, the L-DOPA production with NaAsc under initial conditions requires ∼30 min to surpass that achieved with H_2_O_2_ (Fig. 1C). To further improve the catalytic efficiency of the TyrH/NaAsc system, we systematically optimized key reaction components: NaAsc concentration, HCl used for substrate dissolution, substrate L-Tyr concentration, and TyrH enzyme concentration.

As shown in Fig. 4A, L-DOPA yield increased with higher NaAsc concentrations, reaching a substantial level at 100 mM. Instead, higher HCl concentrations leads to the decreased reaction rates and yields (Fig. 4B). Thus, HCl should be used at the minimum for the dissolution of the substrate (e.g., 0.2 M HCl for 100 mM L-Tyr). In terms of the L-Tyr substrate, the optimal concentration was found to be 1∼ mM. While the lower substrate concentration can reduce the product yield (Fig. 4C), the higher concentrations may lead to the substrate inhibition of catalysis. For the enzyme concentration, the catalytic yield nearly plateaued at 40 µM TyrH, with no obvious increase at 60 µM (Fig. 4D). In summary, optimal reaction conditions were determined as 40 µM TyrH, 1 mM L-Tyr (final HCl concentration 2 mM), and 100 mM NaAsc.

**Fig. 4.**
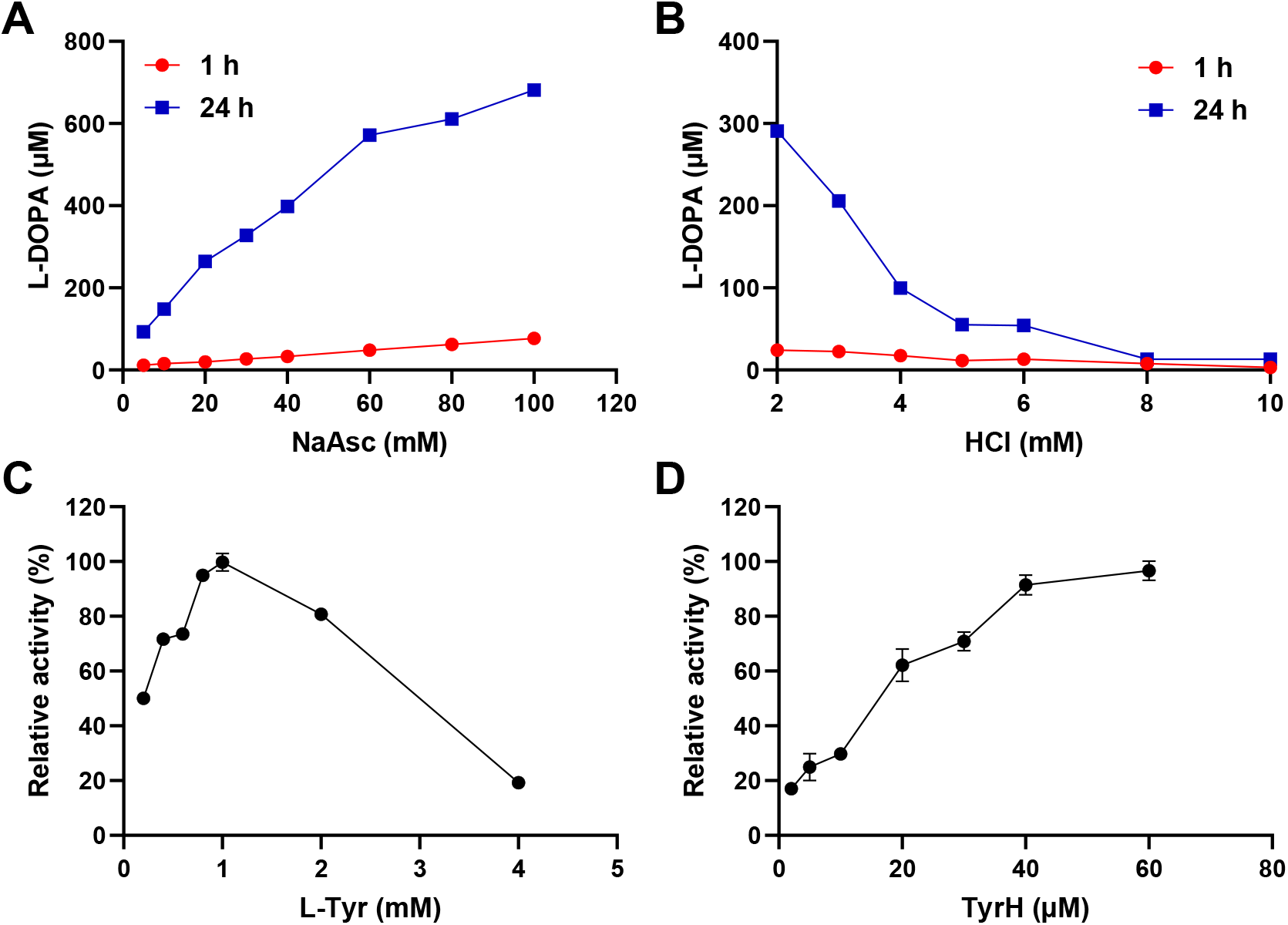
The optimal concentrations of key components of TyrH-mediated hydroxylation. The yield of L-DOPA produced by TyrH under different concentration of NaAsc (A) or HCl (B) on 1 hour and 24 hours. The effect of L-Tyr (C) or TyrH (D) concentration in TyrH-mediated hydroxylation on 4 hours. The product concentration was determined and expressed as relative activity (normalized to the maximum yield obtained). The standard reaction system contained 50 μM TyrH, 1 mM L-Tyr, and 20 mM NaAsc at 30 °C in 50 mM Tris-HCl buffer, pH 7.5, 150 mM NaCl.

Temperature and pH are also critical for enzymatic activity. The highest catalytic rate (after 1 h) and final yield (after 24 h) were achieved at 40 °C (Fig. 5A). Lower temperatures reduced catalytic thermodynamics, whereas temperatures ≥50 °C may lead to the rapid enzyme inactivation and lower product yield. Buffer pH studies showed that the initial rate increased with higher pH, but the final yield after 24 h peaked at pH 8.5, surpassing yields at pH 9.0 and 10.0 (Fig. 5B). Thus, moderately alkaline conditions are beneficial for maintaining TyrH activity. In sum, the optimized temperature and pH were determined to be 40 °C and pH 8.5, respectively.

**Fig. 5.**
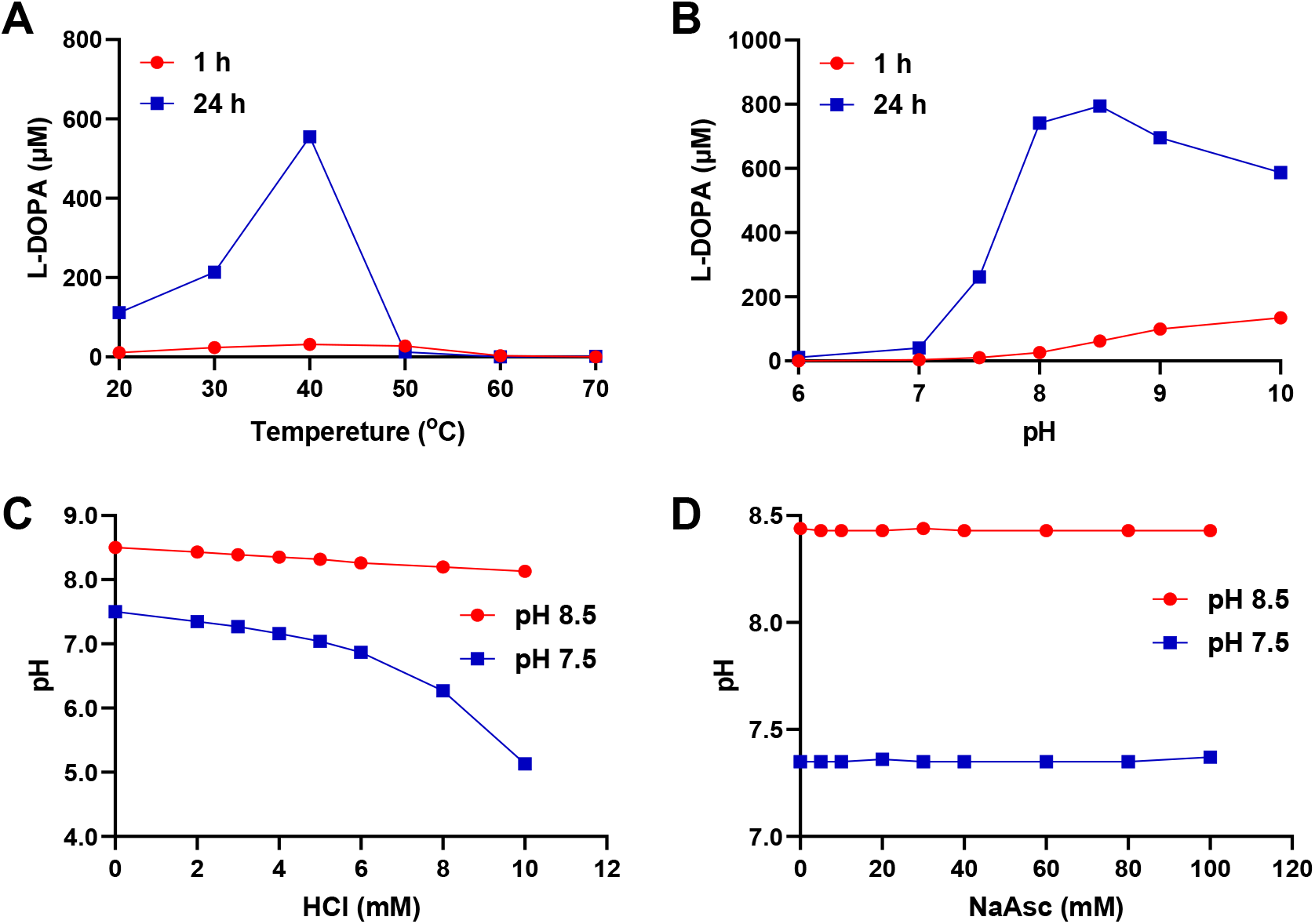
The optimal temperature and pH of TyrH-mediated hydroxylation. (A) The yield of L-DOPA produced by TyrH at various temperatures in buffers of pH 7.5. (B) The yield of L-DOPA produced by TyrH at 30 °C in buffers between pH 6 to 10. The standard reaction system contained 50 μM TyrH, 1 mM L-Tyr, and 20 mM NaAsc at 30 °C in 50 mM Tris-HCl buffer, pH 7.5, 150 mM NaCl. (C) The effect of pH in the reaction system by adding different concentrations HCl. (D) The effect of pH in the reaction system by adding different concentrations NaAsc.

Considering that the addition of L-Tyr (dissolved in 0.2 M HCl) and NaAsc may affect the pH of the reaction system, we measured the buffer pH after their addition. As L-Tyr concentration increases, the pH of a pH 7.5 buffer drops rapidly, even to pH 5.0, whereas a pH 8.5 buffer remained stable, maintaining pH > 8.0 even with a final HCl concentration of 10 mM (Fig. 5C). This confirms that a pH 8.5 buffer better preserves the catalytic reaction pH. Moreover, increasing NaAsc concentration did not alter the pH of either buffer (Fig. 5D), confirming that NaAsc does not affect reaction pH—an advantage for catalytic performance.

### Catalytic Performance of TyrH using NaAsc under Optimized Conditions

Using the optimized conditions (40 µM TyrH, 1 mM L-Tyr, 100 mM NaAsc, 40 °C, pH 8.5), we performed TyrH-mediated hydroxylation of L-Tyr and monitored product formation and substrate consumption. The reaction initially showed a high rate, achieving 70% conversion within the first hour. The rate subsequently decreased, and the reaction was nearly complete after 2 h, reaching a final yield of 95% with almost quantitative substrate conversion (Fig. 6A). This catalytic rate and final yield are substantially higher than under non-optimized conditions (Fig. 1D).

**Fig. 6.**
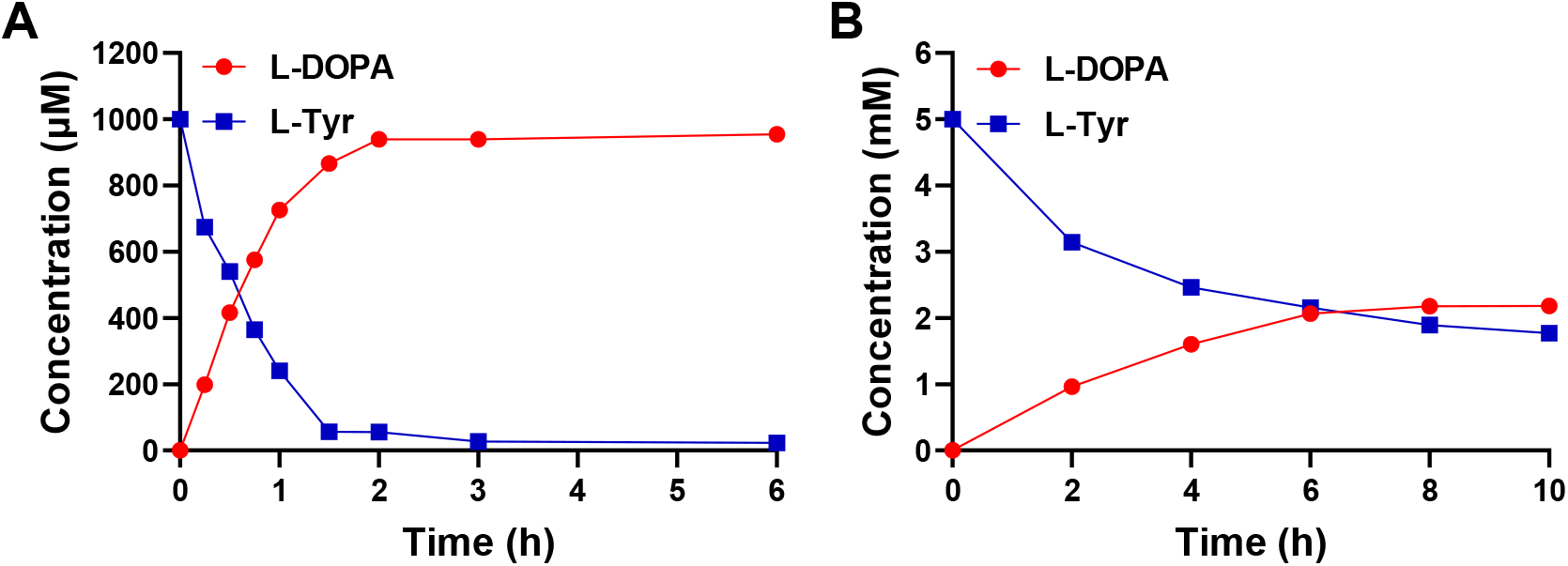
TyrH-mediated hydroxylation of L-Tyr using NaAsc under optimized catalytic conditions. (A) Time course of TyrH-mediated hydroxylation of L-Tyr to L-DOPA using NaAsc as reductant. Reaction was performed in the optimized catalytic conditions (40 µM TyrH, 1 mM L-Tyr, 100 mM NaAsc, 40 °C, pH 8.5). (B) Time course of TyrH-mediated hydroxylation in the optimized catalytic conditions with a high substrate concentration (5 mM L-Tyr).

To assess industrial potential, we investigated catalytic performance at elevated substrate concentrations (Fig. 6B). With 5 mM L-Tyr, ∼1 mM L-DOPA was generated after 2 h, comparable to the yield under optimal low-concentration conditions (Fig. 6A). However, the reaction rate gradually declined upon prolonged incubation, nearly ceasing after 6 h with only 2 mM L-DOPA (40% yield). Extending the reaction to 10 h increased the yield only marginally to 44%.

The fact that high L-Tyr concentrations exert a negligible effect on the pH of the pH 8.5 buffer system (Fig. 5C) rules out pH instability as a cause for the limited yields. A more plausible explanation is substrate inhibition. During normal catalysis, substrate diffuses through a tunnel to access the active site, and subsequent product release resets the enzyme. At high substrate concentrations, we hypothesize that “tunnel congestion”^49^ occurs as incoming substrate molecules encounter product yet to be released, which impairs catalytic turnover. Future efforts to enhance catalytic efficiency at high substrate concentrations should therefore focus on rationally modifying the tunnel bottleneck.

## Conclusions

In summary, this study establishes an efficient and operationally stable ascorbate/O_2_-dependent catalytic route for the synthesis of L-DOPA using heme-dependent tyrosine hydroxylase (TyrH). Compared to the conventional H_2_O_2_-driven system, sodium ascorbate (NaAsc) as a reductant confers superior catalytic durability, enabling sustained L-DOPA production and surpassing the H_2_O_2_-mediated yield within 30 min. Spectroscopic analysis directly correlates the rapid inactivation of heme degradation in the presence of H_2_O_2_, while NaAsc maintains heme integrity. QM/MM simulations provide mechanistic insight for the TyrH/Acs/O_2_ system, indicating that *in situ* generated H_2_O_2_ leads to the active species of Compound I for tyrosine hydroxylation, while avoiding the heme degradation by additional H_2_O_2_ molecules.^43, 50-52^

Through systematic optimization, we identified the reaction conditions (40 µM TyrH, 1 mM L-Tyr, 100 mM NaAsc, 40 °C, pH 8.5) that achieve >95% conversion of L-Tyr to L-DOPA within 2 h, demonstrating a combination of high stability and significantly enhanced catalytic efficiency of TyrH/NaAsc/O_2_ system. These findings not only underscore the general applicability and sustained catalytic capacity of the ascorbate/O_2_ pathway in heme proteins, but also provides a practical biocatalytic strategy for sustainable L-DOPA production.

## Author contributions

B.W. conceived the project and directed the research. L.L. and Z.B. designed and performed experiments. Z.J. undertook the QM/MM study and MD simulations. B.W., L.L. and Z.J. co-wrote the manuscript. A.L. made a critical reading of the manuscript.

## Data availability

The authors declare that all data supporting the findings of this study are available within the article and ESI.

## Acknowledgements

This work has been supported by Scientific Research Innovation Capability Support Project for Young Faculty (ZYGXONJSKYCXNLZCXM-B6), National Natural Science Foundation of China (22121001), and Fundamental Research Funds for the Central Universities (20720240124).

