## Supporting Information for "An efficient and stable ascorbate/O_2_‑driven route for L‑DOPA synthesis by heme‑dependent tyrosine hydroxylase"

### 1. Materials and methods

#### 1.1 Materials

LB medium, 20X PBS Buffer, TureColor Marker and Modified Bradford Protein Assay Kit were obtained from Sangon Biotech. BL21(DE3) competent cells was provided by Weidibio. Kanamycin and SDS-PAGE were supplied by Solarbio. Tris, isopropyl ß-D-1-thiogalactopyranoside and imidazole were procured from Acmec. δ-aminolevulinic acid, L-Tyr, L-DOPA, ascorbate and sodium ascorbate were provided by Energy Chemical. Trifluoroacetic acid and acetonitrile were acquired from Sigma-Aldrich. Other chemicals and reagents were the products of Sinopharm.

#### 1.2 Cloning, expression and purification of TyrH

The gene encoding TyrH from *Streptomyces sclerotialus* (GenBank WP_051872337, PDB ID: 7KQR)[1] was synthetized after codon optimization for *E. coli* (GenScript, Nanjing, China) and inserted into the vector pET-28a(+) with the N-terminal $His_6$-tag. The pET-28a(+)-TyrH plasmid was transformed into *E. coli* BL21(DE3) competent cells for protein production. Transformants were grown at 37 °C in LB medium supplemented with 40 mg/L kanamycin. Protein expression was induced at an $OD_{600}$ of 0.6–0.8, with induction performed as follows: 0.5 mM isopropyl ß-D-1-thiogalactopyranoside (IPTG) and 0.3 mM δ-aminolevulinic acid (5-ALA), followed by 20 h of incubation at 30 °C.

The AKTAxpress system (GE healthcare) was used for Purification of protein.[2] The supernatants were was added to 5 mL His-Trap column (GE Healthcare) which pre-equilibrated with 20 mM $Na_3PO_4$, 500 mM NaCl, and 20 mM imidazole, pH 7.4 (binding buffer), and the column was equilibrated by binding buffer. The proteins were eluted by eluting buffer (20 mM $Na_3PO_4$, 500 mM NaCl, and 500 mM imidazole, pH 7.4). Target proteins were desalted by loading onto a PD-10 column (PD SpinTrap G-25, Cytiva) and buffer exchanged in 50 mM Tris-HCl buffer, pH 7.5, 150 mM NaCl. The protein concentrations were determined using a modified Bradford Protein Assay Kit and analyzed by SDS-PAGE. The proteins were concentrated with a 10 kD Ultrafiltration centrifugal filters (Merck Millipore).

UV−vis absorption spectra of TyrH (10−20 μM) were recorded between 300 and 700 nm in 50 mM Tris-HCl buffer (pH 7.5) containing 150 mM NaCl. The concentration of TyrH was determined spectrophotometrically using the Soret band extinction coefficient of horseradish peroxidase (HRP) ($\varepsilon_{Soret}$ = 102 $mM^{-1}$ $cm^{-1}$),[3] based on the structural similarity of their His-ligated heme centers and corresponding Soret absorbance profiles.

**1.3 Heme Bleaching Assay**

Heme bleaching experiments were performed according to the previously published procedure.[4] UV-vis spectroscopy was performed at 30 °C using a microplate reader (Thermo Fisher Scientific). The data interval was l.0 nm, and the data range was 700 to 300 nm. A baseline spectrum was first recorded with 200 µL of 20 µM TyrH in 50 mM Tris-HCl, pH 7.5, 150 mM NaCl. The different concentration of $H_2O_2$ or NaAsc (5, 10, 20 mM) were added to TyrH and mixed for 10 min. Subsequent, the UV-vis spectrum was recorded to monitor the heme destruction (bleaching of the Soret band) in different cofactors. Meanwhile, $H_2O_2$ (1 mM) or NaAsc (20 mM) was added, and spectra were recorded at different time to monitor the rate of heme destruction.

**1.4 The activity assay of TyrH**

The standard reaction system contained 20 µM TyrH, 1 mM L-Tyr (from a 100 mM stock solution dissolved in 0.2 M HCl), and 1 mM $H_2O_2$ or 20 mM NaAsc/Asc in 50 mM Tris-HCl buffer, pH 7.5, 150 mM NaCl. Reactions with TyrH were conducted at 30 °C and 900 rpm for 24 h. a 50 µL aliquot of the reaction mixture was mixed with 2.5 µL of concentrated HCl (6 M) to quench the reaction at different time. Then 152 µL of water was added to obtain a 200 µL final volume of sample.

Product analysis of TyrH was performed using a HPLC system (Waters) equipped with a ZORBAX-Eclipse Plus C18 column (250 mm × 4.6 mm, 5 µm; Agilent). Mobile phases consisted of 0.1% (v/v) trifluoroacetic acid in water (solvent A) and methanol (solvent B). The program was as follows: 0–10 min, 10% B, at a flow rate of 1 mL/min. Detection was performed at 280 nm.

**1.5 The optimal reaction concentration on TyrH reaction**

The standard reaction mixture consisted of 50 µM TyrH, 1 mM L-Tyr, and 20 mM NaAsc in 50 mM Tris-HCl buffer (pH 7.5) containing 150 mM NaCl. Reactions were incubated at 30 °C for 24 h. To determine the optimal concentration of NaAsc, reactions were performed with varying

NaAsc levels (5, 10, 20, 30, 40, 60, 80, and 100 mM) while keeping all other components as described in the standard system. Product formation was measured after 1 h (initial rate) and 24 h (final yield). Because L-Tyr was dissolved in HCl, the final HCl concentration in the reaction could influence the pH of the assay buffer. To evaluate this, the final HCl concentration was varied (2, 3, 4, 5, 6, 8, and 10 mM) while maintaining 1 mM L-Tyr and other standard conditions. The initial rate (1 h) and final yield (24 h) were determined.

The effects of substrate and enzyme concentration on catalytic performance were also systematically evaluated. The effect of L-Tyr concentration on TyrH reaction was tested by varying the final L-Tyr concentration (0.2, 0.4, 0.6, 0.8, 1.0, 2.0, and 4.0 mM) under otherwise standard conditions. Reactions were stopped after 4 h, and the product concentration was measured. Relative activity was calculated by normalizing all values to the maximum product yield (set as 100%). To assess the influence of enzyme loading, TyrH concentration was varied (2, 5, 10, 20, 30, 40, and 60 μM) while keeping other components constant. After 4 h of reaction, the product concentration was determined and expressed as relative activity (normalized to the maximum yield obtained).

#### 1.6 The optimal temperature and pH investigation on TyrH reaction

The influence of reaction temperature and buffer pH on enzyme-catalyzed reactions is critical. Therefore, we examined the effects of temperature and buffer pH on the TyrH reaction. To assess the impact of temperature, the standard reaction system (as described previously) was used while varying the incubation temperature (20, 30, 40, 50, 60, and 70 °C). The initial reaction rate (after 1 h) and final product yield (after 24 h) were determined under otherwise identical conditions. To evaluate the effect of buffer pH, the standard reaction was carried out in 50 mM Tris-HCl buffer (pH 6.0, 7.0, 7.5, 8.5, 9.0, or 10.0) containing 150 mM NaCl, while keeping all other components unchanged. The initial rate (1 h) and final yield (24 h) were also measured.

Additionally, to analyze how the addition of HCl (used for substrate solubilization) and the cofactor NaAsc might alter the pH of the reaction buffer, the following pH measurements were conducted: To 50 mM Tris-HCl buffer (pH 7.5 or 8.5) containing 150 mM NaCl and 20 mM NaAsc, different final concentrations of HCl (0, 2, 3, 4, 5, 6, 8, and 10 mM) were added, and the resulting pH was recorded using a calibrated pH meter; To 50 mM Tris-HCl buffer (pH 7.5 or 8.5) containing

150 mM NaCl and 1 mM L-Tyr (with a final HCl concentration of 2 mM from the substrate stock), different final concentrations of NaAsc (0, 5, 10, 20, 30, 40, 60, 80, and 100 mM) were added, and the pH was measured accordingly.

**1.7 The optimal condition on TyrH reaction**

Using the optimized catalytic conditions (40 μM TyrH, 1 mM L-Tyr, 100 mM NaAsc, 40 °C, pH 8.5), the TyrH-catalyzed reaction was performed and monitored over time. Aliquots were withdrawn at specific time points (15, 30, 45, 60, 90, 120, 180, and 360 min), and the concentrations of the product formed and the remaining substrate were analyzed by HPLC, as described in the previous method. To evaluate the catalytic capacity of TyrH under high substrate loading, the reaction was also carried out with an increased L-Tyr concentration of 5 mM (resulting in a final HCl concentration of 10 mM) while keeping all other conditions unchanged. Samples were taken at 2, 4, 6, 8, and 10 h, and similarly analyzed by HPLC to determine product formation and substrate depletion.

**1.8 System setup and molecular dynamics simulations**

The initial structure was obtained from the crystal structure of TyrH (PDB entry code: 7KQR, solved at 1.89 Å resolution).[1] Protonation states of titratable residues (Glu, Asp, His) were determined via PROPKA[5] analysis combined with visual inspection of hydrogen-bonding networks. Partial atomic charges for substrates were derived using restrained electrostatic potential (RESP) calculations[6] at the HF/6-31G* level[7, 8]. The iron ion and its coordination sphere were parametrized using MCPB.[9, 10] To neutralize the total charges of the systems, some $Na^+$ ions were added to the proteins' surfaces. Additionally, the systems were dissolved in a cube of TIP3P waters with a 10 Å water layer. The protein residues were described by the Amber ff19SB force field[11] and general AMBER force field (GAFF) [12]. Topology files and initial coordinates were prepared using the tleap module of AMBER 18.[13-15] After the system setup was complete, it was minimized by a combined steepest descent and conjugate gradient method. Then the system was heated from 0 to 300 K for 50 ps with 0.5fs time step under a constant pressure. Afterwards, 1 ns NPT equilibration was used to relax the density of system to 1.0 $g\cdot cm^{-3}$ and 4ns further equilibrated the system without all restraints under NPT ensemble to get an equilibrated pressure and temperature. At last, 100ns NPT MD simulations were conducted at 300K. All covalent bonds involving hydrogen atoms were constrained using SHAKE algorithm.[16]

#### 1.9 QM/MM Calculations

The representative snapshots extracted from MD trajectories were used for the subsequent QM/MM calculations. All the QM/MM calculations were conducted using ChemShell,[17-19] which combines Turbomole[3] for the QM region and DL_POLY[20] for the MM region. The polarizing effect of the enzyme environment on the QM region was treated via an electrostatic embedding scheme. [21] The QM region includes the L-Ascorbate, truncated Heme, the axial His ligand, the truncated His88, and key waters in the active site. The QM part of the system was treated by TPSSH hybrid *meta*-GGA functional[22] with the all-electron basis set of def2-SVP[23], whereas the MM part was modelled at the classical level using the same parameters as in the molecular dynamics simulations. Energies were further corrected with the large all-electron basis set def2-TZVP, supplemented by Grimme's D3 dispersion correction.[24-26] Transition states were found with relaxed potential energy surface scans and then optimized by dimer optimizer implemented in DL-FIND code.[17]

### 2. Supplementary figure and table

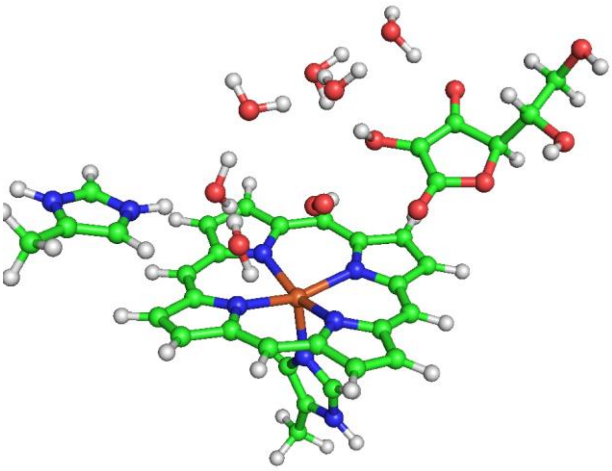

**Figure S1.** The QM region used for TyrH/ASC/$O_2$ system.

**Table S1.** Spin density population of key atoms for key species involved in the TyrH/ASC/$O_2$ system.

| Species | Spin density | | | | | |
|---|---|---|---|---|---|---|
| | Fe | Op | Od | His-88 | HEM | ASC |
| $^8$RC | 4.27 | 1.01 | 0.96 | 0.10 | 0.63 | 0.00 |
| $^2$IM1 | 1.19 | -0.51 | -0.53 | -0.03 | 0.01 | 0.90 |
| $^2$IM2 | 1.10 | -0.09 | -0.08 | 0.02 | -0.73 | 0.77 |
| $^2$PC | 1.07 | 0.00 | 0.00 | 0.00 | -0.70 | 0.63 |

### 3. Amino acid sequence

**Amino acid sequence of TyrH:**

MNTGTGTVLTELPDHGRWDFGDFPYGLEPLTLPEPGSLEAADSGSVPAEFTLTCRHIAAIA
AGGGPAERVQPADSSDRLYWFRWITGHQVTFILWQLLSRELARLPEEGPERDAALKAMTR
YVRGYCAMLLYTGSMPRTVYGDVIRPSMFLQHPGFSGTWAPDHKPVQALFRGKKLPCVR
DSADLAQAVHVYQVIHAGIAARMVPSGRSLLQEASVPSGVQHPDVLGVVYDNYFLTLRS
RPSSRDVVAQLLRRLTAIALDVKDNALYPDGREAGSELPEELTRPEVTGHERDFLAILSEVA
EEATGSPALASDR
